# Developing permeable polydimethylsiloxane-based biomimetic leaf surfaces to study phyllosphere microbial ecology

**DOI:** 10.64898/2026.05.12.724494

**Authors:** Evan Kear, Michał Bernach, Volker Nock, Mitja Remus-Emsermann

## Abstract

Polydimethylsiloxane (PDMS) is an excellent material for the construction of biomimetic leaf replicas which reproduce leaf surfaces with high fidelity. This allows for the study of leaf surface-colonizing bacteria and the impact of the leaf topology on bacterial distributions and behavior. However, their application is limited to short-term experiments, as long term survival of microorganisms on their surface is not possible due to a lack of nutrient replenishment. On living leaves, nutrients diffuse across the cuticle via leaching, a process not yet replicated in biomimetic systems. Here, we explore whether water and fructose can be supplied to microbial colonizers on PDMS membranes by mimicking leaching. We created hybrid membranes by incorporating polymers (Carbopol, Pemulen, cellulose microfibers, cellulose nanocrystals, and polyvinylpyrrolidone) to enhance nutrient transport. We determined that bulk diffusion of water correlated negatively with membrane thickness and positively with polymer concentration. Further, fructose diffusion across hybrid membranes reached similar rates compared to isolated *Populus × canescens* leaf cuticles.

Under high relative humidity, these membranes supported long-term bacterial survival. Our findings represent important steps towards the development of topomimetic leaf surfaces that sustain microbial life, enabling further investigation into the microbe-microbe interactions that take place on leaves.

## Introduction

Leaf-associated microorganisms exhibit high diversity and non-random, heterogeneous spatial distributions [1,2]. The most prevalent group of microorganisms on leaves are bacteria [3]. The first layer of contact between those microorganisms and the plant is the leaf cuticle which is composed of cutin and cuticular waxes [4,5]. The cuticle is usually highly hydrophobic and thereby determines the interactions with the environment, protecting the plant from pathogen ingress and preventing the loss of water and solutes [6]. Despite its role in prevention of water and solute loss, a noteworthy amount of nutrients, including carbohydrates, amino acids and organic acids, diffuses from the leaf interior through the process of leaching [7,8]. Leaching does not take place homogeneously across the cuticle; instead, leaching results in nutrient rich sites on the leaf surface, which can be accessed by microorganisms [9,10]. Often those sites also co-occur with the phyllothelma - the waterbodies present on leaf surfaces [11]. Several studies have demonstrated heterogeneous distribution of nutrients, such as iron or fructose, on plant leaf surfaces or isolated leaf cuticles [12,13]. The heterogeneity of the available resources leads to microhabitats that feature different levels of habitability and are able to sustain different levels of colonization [14]. Further heterogeneity is introduced by the structure of the leaf surface, with features such as epidermal grooves, trichomes, and stomata contributing to a complex topographical landscape. These features can generate preferential environments for bacterial colonization through protection from abiotic stressors, or through increasing nutrient availability [2,13]. It is also possible that leaf topography alone modulates bacterial interactions, through processes such as limiting secondary metabolism, preventing aggregation, or forcing cooperative behaviors [15–17]. Although these mechanisms have been observed prior on spatially complex surfaces, they have not been observed on the leaf surface. More factors, such as colonization sequence or local microbial community composition, can additionally affect the fate of individual microorganisms [2,18,19].

It has been well established that elastomeric polydimethylsiloxane (PDMS) is an ideal compound for the fabrication of biomimetic leaf surfaces [20]. PDMS has been used to produce a number of biomimetic leaf surfaces including *Spinacia oleracea* L. [21–23], *A. thaliana* [24], *Phaseolus vulgaris* L., *Lactuca sativa* L., *Solanum lycopersicum* L., *S. tuberosum* L., *C. zebrina* [11], *Lactuca sativa, Lactuca sativa* L. var. *longifolia*, *Brassica oleracea* [23]. Although extremely useful to study several aspects of bacterial life on leaves — such as the impact of leaf topography on bacterial survival after disinfection [22,23] or the distribution of water on leaves [11] — these PDMS leaf replicas do not sustain microbial life because nutrient availability is limited and not replenished post-inoculation, thereby limiting investigation involving community interactions or dynamics.

In this study, we investigated the effect of several filler polymers—Carbopol^®^, Pemulen^™^ (from our previous work [25]), as well as cellulose microfibers (CMFs), cellulose nanocrystals (CNCs), and polyvinylpyrrolidone (PVP)—on nutrient diffusion (leaching) to the surface of PDMS-based membranes. Carbopol, Pemulen, and PVP were selected as hygroscopic additives to increase surface water retention on the otherwise hydrophobic PDMS, thereby promoting the diffusion and availability of hydrophilic compounds [26,27].

CMFs were included due to their natural occurrence in leaf cuticles, where they contribute to the formation of polar pathways which facilitate solute transport [28]. CNCs were incorporated for their reported ability to generate asymmetric diffusion barriers that mimic those of leaf surfaces [29]. Together, these additives were hypothesized to create hydrated, permeable networks within the PDMS matrix, enhancing the transport of larger polar molecules across the membrane. As a first step, we assessed the bulk diffusion of water across both pure and hybrid PDMS membranes to evaluate their baseline permeability.

Subsequently, we measured the permeability of the membranes to fructose, a major photosynthate commonly found on leaf surfaces [13]. Fructose bulk diffusion was quantified using a fructose-responsive fluorescent whole-cell bacterial bioreporter based on the model leaf surface colonizer *Pantoea eucalypti* 299R (formerly *Pantoea agglomerans* 299R or *Erwinia amylovora* 299R) [10,13,30]. Finally, we assessed the survival of fluorescent *P. eucalypti* 299R on PDMS and hybrid PDMS membranes under varying nutrient and humidity conditions.

## Results

### Water permeability of PDMS and hybrid PDMS

We first assessed membrane thickness effects at constant filler polymer concentration (10%). In all cases, thinner membranes were more permeable to water (Figure 1 A). The normalized bulk diffusion measured using the inverted diffusion chamber system (PDMS + 10% CNC treatment) was a magnitude higher compared to the steel diffusion chamber system (all other treatments). The discrepancy between diffusion values between experimental systems is thought to arise due to increased fluid pressure and subsequent bulging of the membrane, thus increasing the surface area for diffusion while simultaneously lowering the true thickness of the membrane. This effect is corrected by normalizing the data using the PDMS controls from each system, generating a relative flux value (Figure 1 B).

**Figure 1:**
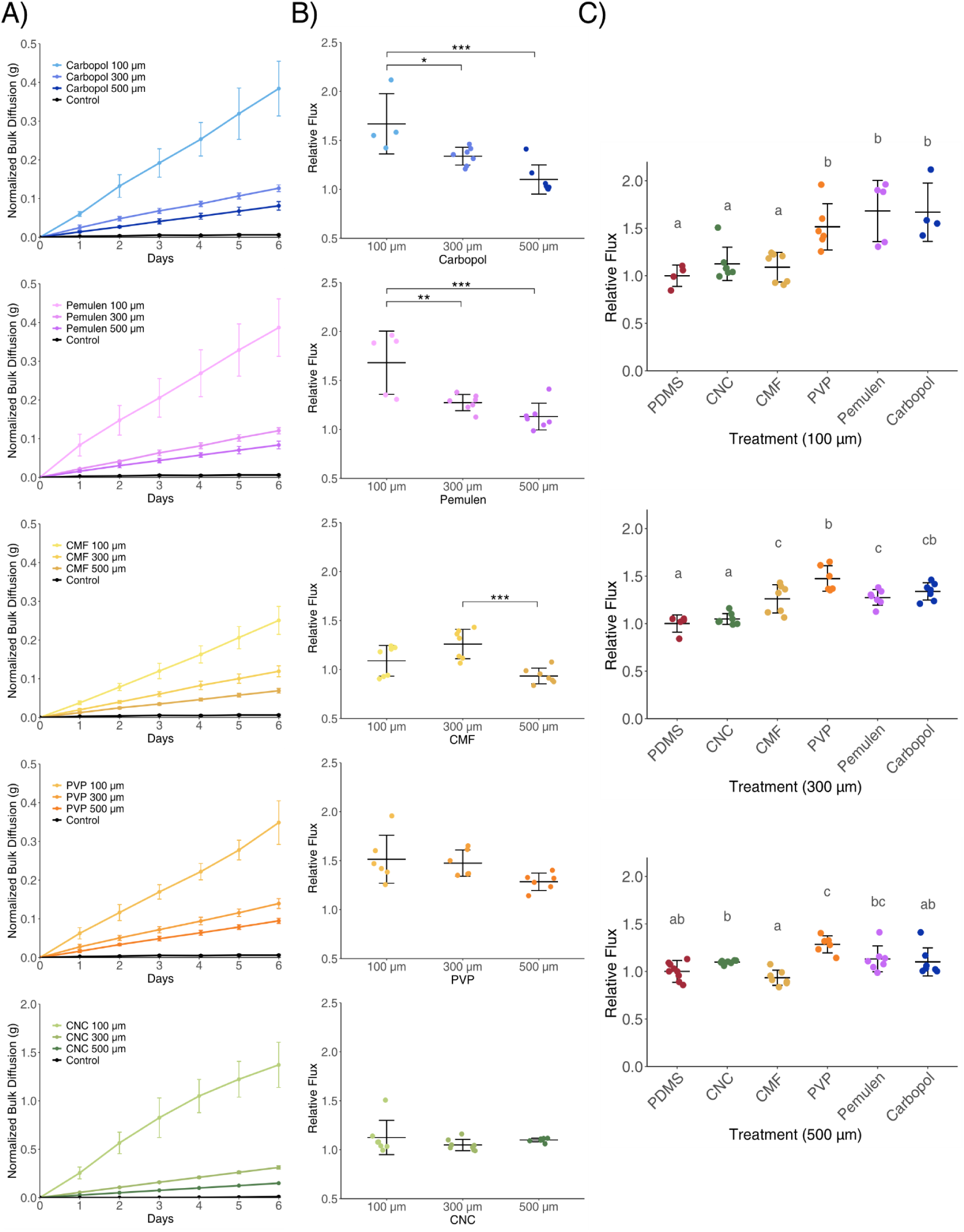
The influence of membrane thickness on water diffusion rates through hybrid PDMS. A) Normalized bulk water diffusion coefficients through 100, 300, and 500 μm hybrid PDMS membranes with 10% Carbopol, Pemulen, CMF, PVP, or CNC. B) Water flux relative to thickness-matched PDMS controls. Asterisks denote Tukey’s post-hoc p-values (p=0.05, p=0.01, p=0.001). C) Pairwise relative flux comparisons between pure PDMS and 10% hybrids per thickness Treatments with the same letter(s) are not significantly different (Tukey’s post-hoc test, p=0.05. n = 7). Points represent single membranes and error bars denote standard deviation.

This normalization also removes the effect of membrane thickness. Within each filler polymer treatment, thinner PDMS + 10% Carbopol and PDMS + 10% Pemulen membranes showed higher relative flux than thicker ones, revealing a thickness-filler polymer interaction; PDMS + 10% PVP and PDMS + 10% CNC did not show this interaction (no significant flux differences across thicknesses). Relative to pure PDMS controls (Supplementary Figure 3), flux through PDMS + 10% PVP was significantly higher at all thicknesses (100 μm, P=0.01621; 300 μm, P<0.001; 500 μm, P<0.001). PDMS + 10% Pemulen and PDMS + 10% Carbopol were higher at 100 μm and 300 μm (but not 500 μm). PDMS + 10% CMF was higher only at 300 μm. PDMS + 10% CNC showed no significant differences at any thickness.

Next, we determined the effect of filler polymer concentration at a constant membrane thickness of 300 μm which showed that higher filler polymer concentrations generally increased water permeability (Figure 2A). The inverted diffusion chamber setup consistently showed higher permeability across all concentrations for PDMS + CNC membranes, which the normalization process accounted for. Notably, 15% w/w filler polymer concentrations yielded significantly higher permeability than pure PDMS for all filler polymers, whereas only PDMS + PVP and PDMS + Carbopol showed increases at 5% w/w.

**Figure 2:**
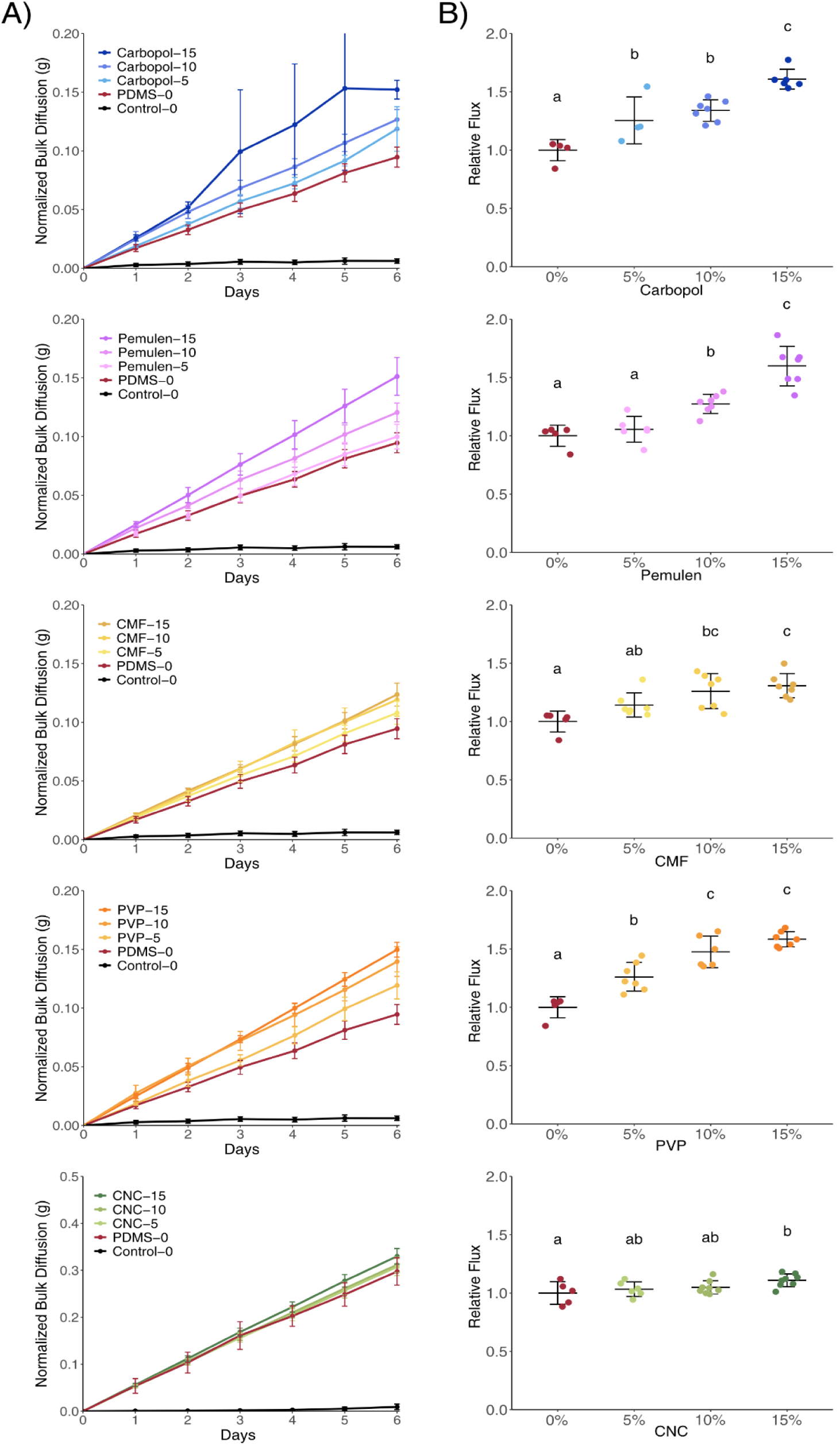
The influence of filler concentration on water diffusion rates through PDMS and hybrid PDMS. A) Water diffusion through membranes composed of 300 μm PDMS, 5, 10 and 15% PDMS + Carbopol, Pemulen, CMF or PVP. Generally, higher concentrations of fillers lead to higher diffusion. B) Pairwise comparisons of relative flux between 300 μm hybrid PDMS membranes at different filler concentrations. Treatments with the same letter(s) are not significantly different (Tukey’s post-hoc test, p=0.05, n = 7). Points represent single membranes and error bars denote standard deviation.

### Fructose permeability of PDMS and hybrid PDMS

To investigate whether the addition of filler polymers increased PDMS membrane permeability to fructose, Carbopol, Pemulen, CMF, CNCs and PVP were mixed into the polymer to the concentration of 10% w/w. For each material, 300 μm thick membranes were fabricated. Bulk fructose diffusion was determined using diffusion chambers [39]. The medium in the donor chamber included 3 M fructose and samples were collected after 6 and 12 days. Following sample collection, the concentration of fructose in medium from the acceptor chamber was determined using the fluorescent whole-cell bacterial bioreporter Pe299FruByfp. Experiments were controlled using known concentrations of fructose paired with the fitting of a Monod model (Figure 3 A and B). Compared to pure PDMS (1.82 × 10⁻^9^ kg m⁻^2^ s⁻^1^), the incorporation of 10% Pemulen and 10% Carbopol increased fructose flux by up to four orders of magnitude (1.05 × 10⁻^5^ kg m⁻^2^ s⁻^1^, 8.92 × 10⁻^6^ kg m⁻^2^ s⁻^1^, respectively), while 10% PVP increased flux by approximately three orders of magnitude (1.62 × 10⁻^6^ kg m⁻^2^ s⁻^1^). These diffusion values were similar to the flux observed for isolated *Populus* × *canescens* leaf cuticles (1.69 × 10⁻^6^ kg m⁻^2^ s⁻^1^) [10]. In contrast, 10% CMF (1.94 × 10⁻^9^ kg m⁻^2^ s⁻^1^) and 10% CNC (2.78 × 10⁻^9^ kg m⁻^2^ s⁻^1^) hybrid PDMS membranes did not differ significantly from the pure PDMS membranes or the control treatment (1.70 × 10⁻^9^ kg m⁻^2^ s⁻^1^).

**Figure 3:**
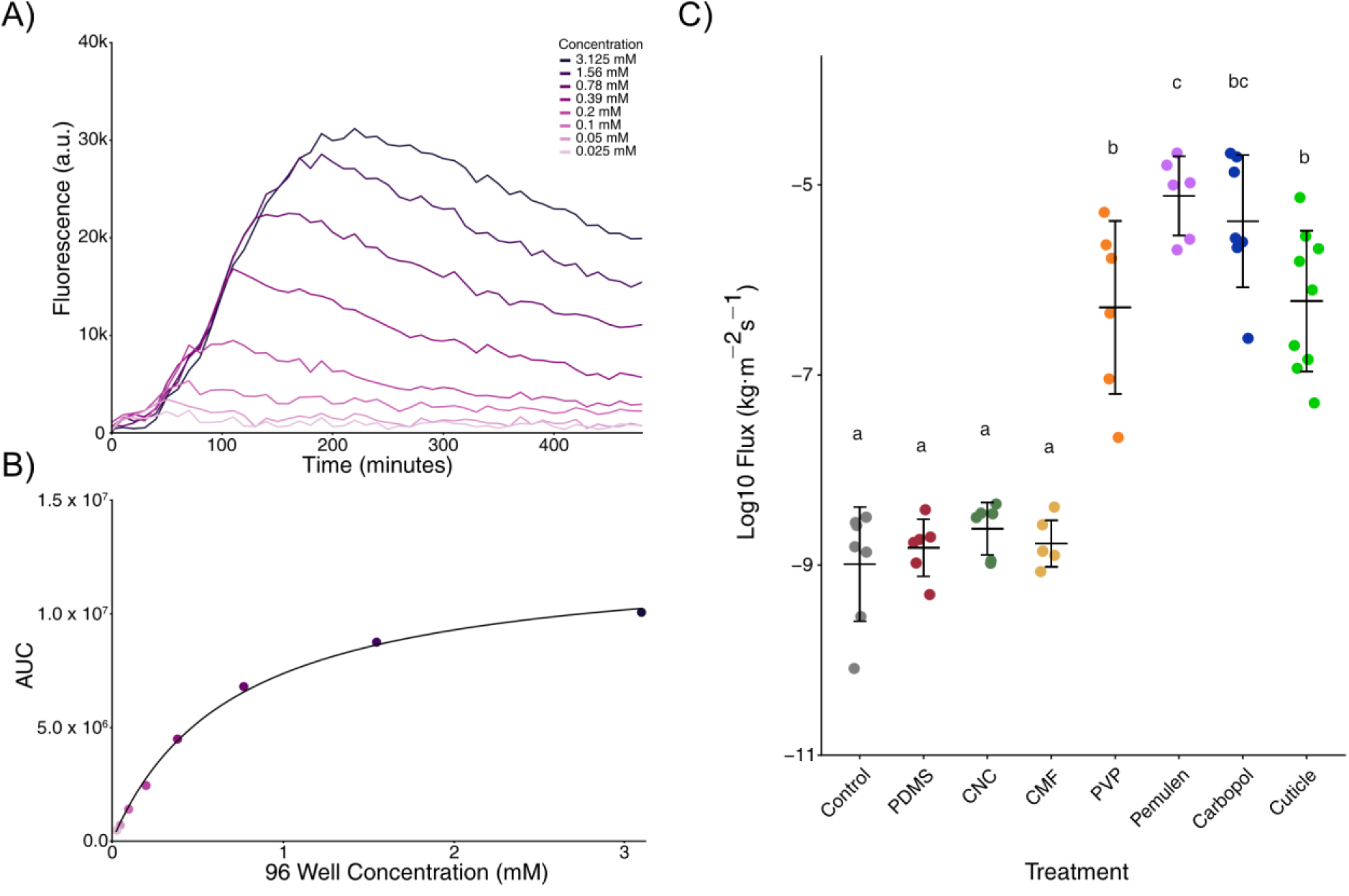
Diffusion of fructose through PDMS and hybrid PDMS. P. eucalypti 299R FruBP fluorescent bioreporter strain enables dose-dependent detection of fructose. Fluorescence was measured over time for samples with known fructose concentrations. Fluorescence over time measured for samples with known concentration of fructose. (A) A standard curve was generated by plotting the area under the curve (AUC) against fructose concentration. (B) Separate standard curves were prepared for each measurement. (C) Fructose concentrations in unknown samples were interpolated from these curves, and flux was subsequently calculated. Flux values for 300 μm PDMS and hybrid PDMS membranes were compared with fructose flux across isolated Populus × canescens leaf cuticles (data from [10]). A 250 μm PVC film served as a negative control. Pairwise comparisons between PDMS and 10% hybrid membranes were performed using Tukey’s post hoc test. Treatments sharing the same letters are not significantly different (p = 0.05, n = 6–8). Points represent single membranes and error bars denote standard deviation.

### Bacterial survival on PDMS and hybrid PDMS

To investigate if fructose permeability of PDMS and hybrid PDMS membranes is sufficient to sustain bacterial life on the membrane surfaces, bacterial survival was assessed. Pemulen was chosen as the sole filler for hybrid PDMS fabrication, as the addition of 10% Pemulen increased permeability of PDMS to fructose the greatest. Survival of Pe299_yfp was assessed after 12 and 24 hours. After 12 hours post inoculation, between 1 and 6.5% of the bacteria were culturable compared to T0. After 24 hours post inoculation, similar population sizes remained culturable (Supplementary Figure 4). There was no significant difference in survival of bacteria between PDMS and hybrid PDMS membranes.

No survival occurred beyond 48 hours at 75% or 85% relative humidity (RH), whereas limited growth was observed at 95% RH (data not shown). Bacteria survived for 12 days on pure PDMS membranes at 100% RH, with culturable numbers decreasing by roughly one order of magnitude from the T0 sample (Figure 4). This observation was only true in the presence of growth media containing fructose, bacteria could not be recovered after 96 hours under the nutrient control media (lacking fructose).

**Figure 4:**
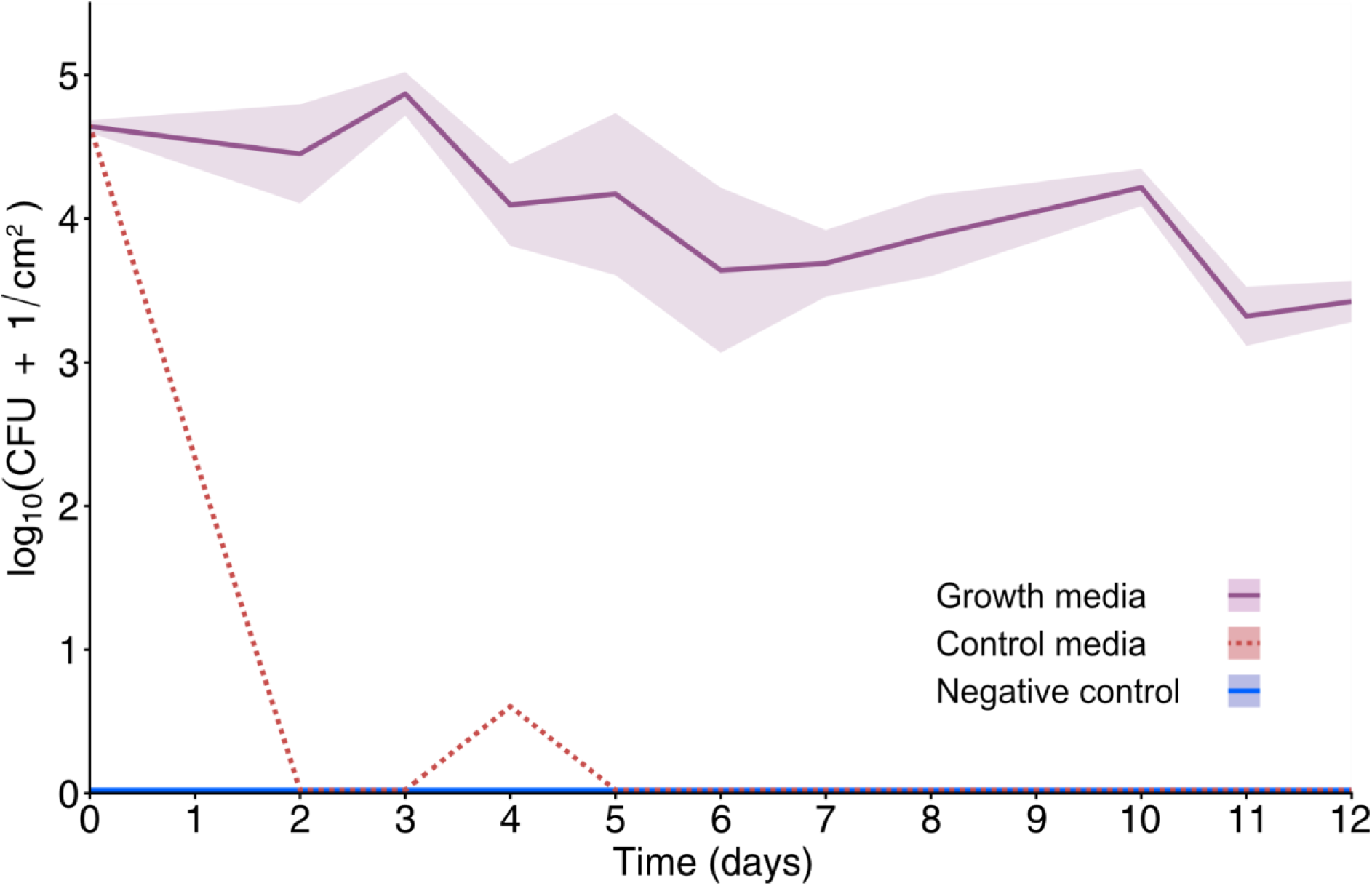
Long Term Survival at 100% RH. *P. eucalypti* 299R was inoculated onto PDMS membranes mounted on the inverted diffusion chamber system and incubated at 100% relative humidity. Membranes were sampled for CFU counts every 24 hours for 12 days. The uninoculated negative control membranes remained sterile throughout the experiment. After 96 hours, single colonies were observed on the membrane located beneath the control medium. By contrast, the growth medium samples maintained high CFU counts, showing a gradual decline over the course of the experiment. (n = 6). Error envelop represents standard deviation.

### Fructose availability on the surface of PDMS and hybrid PDMS

To investigate the distribution of fructose on membrane surfaces Pe299FruByfp_red was used. The red fluorescent signal from Pe299FruByfp_red allows tracking the strain, while the yellow fluorescence indicates the presence of fructose. Bacteria were visualized at 0 and 4 hours post inoculation. Pe299FruByfp_red bioreporter signals were observed on the surface of both PDMS and PDMS + 10% Pemulen membranes (Figure 5). Bacteria were found in sites that originated from droplets after inoculation. On PDMS, those droplets have a sharp, entire edge, while on PDMS + 10% Pemulen, the droplets have ragged edges. No yellow fluorescence was detected at T_0_. After 4 hours, when the membrane was exposed to fructose, fructose-induced bioreporter signals could be detected. This was independent of whether the membranes were supplemented with 10% Pemulen or not. These sites exhibiting yellow fluorescent cells were heterogeneously distributed. Within these sites, the signal is stronger in the center of bacterial aggregates.

**Figure 5:**
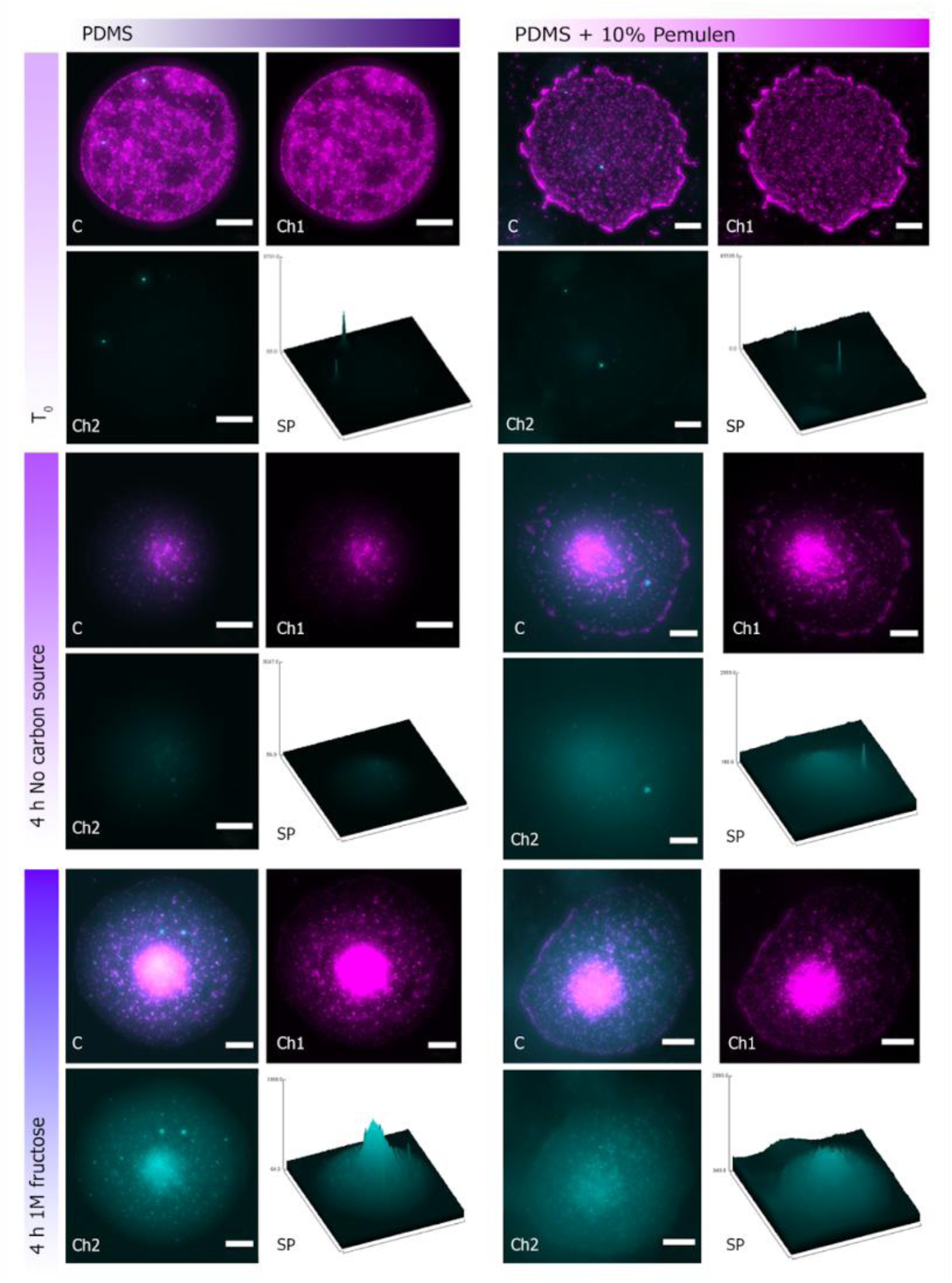
Fructose availability on the surface of PDMS and hybrid PDMS. Pe299FruByfp_red was tracked on the surface of PDMS and PDMS + 10% Pemulen. Red fluorescent signal (magenta) originated from the constitutively expressed mScarlet-I protein. Yellow fluorescent signal (cyan) is the result of fructose inducible production of yellow fluorescent protein. Microsites, where fructose was detected, were present on the surface of both materials when fructose was supplied and were heterogeneously distributed. Scale bar represents 50 μm. ‘C’ in the lower left of an image refers to a composite image, ‘Ch1’ to red fluorescence channel, ‘Ch2’ to yellow fluorescence channel and ‘SP’ to surface plot of the yellow fluorescence.

## Discussion

PDMS has previously been introduced to phyllosphere microbiology to mimic the leaf surface [11,21,22]. However, sustained microbial growth has not yet been demonstrated on PDMS leaf replicas, likely due to the lack of nutrient availability on these surfaces. To address this limitation, we explored modulating the diffusion characteristics of PDMS membranes through the enrichment of PDMS with filler polymers, expanding on our previous progress [25].

5%, 10%, and 15% w/w Carbopol, Pemulen, CMF, CNC, and PVP hybrid PDMS were assessed. These concentrations of fillers were chosen based on previous results that showed filler polymers significantly increased water permeability of 300 μm PDMS membranes [25]. The filler polymers Carbopol and Pemulen were selected for their superabsorber properties, and previous indication of increasing permeabilities through membranes [26]. PVP is also highly hygroscopic and can absorb large quantities of water [27,45]. Cellulose microfibers were chosen since they are naturally occurring in leaf cuticles and very likely involved in shaping the permeability properties of cuticular membranes [28]. CNCs, a variant of cellulose, were chosen for their ability to mimic the asymmetric membrane permeability of the leaf cuticle [29]. The incorporation of these compounds aims to generate permeable networks throughout the PDMS membrane, allowing the diffusion of larger hydrophilic compounds, such as sugars, across the PDMS surface.

As expected, membrane thickness was negatively correlated with bulk water diffusion for all filler polymers. After normalizing data against pure PDMS membranes of the same thickness, the effect of thickness was effectively removed, and mass transport could be compared as relative flux. When relative flux in hybrid PDMS was significantly higher than in pure PDMS, but did not differ significantly between thicknesses, the filler polymer increased permeability by a factor independent of membrane thickness, as observed for PDMS + 10% PVP membranes. By contrast, a significant difference in relative flux after normalization indicates an interaction between filler polymer and membrane thickness, as seen for PDMS + 10% Carbopol and PDMS + 10% Pemulen, where the effect of the filler polymer increased as membrane thickness decreased.

Although incorporating CNCs into PDMS membranes has been reported to produce diffusion properties resembling those of true leaf cuticles [29], in this study significantly higher water diffusion compared to pure PDMS was only observed for the 300 µm PDMS + 15% CNC membranes. One likely explanation is the difference in membrane thickness between studies: Kamtsikakis et al. used membranes as thin as 30 µm, whereas the thinnest membranes here were 100 µm in thickness. At lower thicknesses, CNCs may bridge the membrane, creating continuous diffusion pathways. While individual CNC particles are too small to span the membrane thickness (length ∼150 nm), CNC powder has a particle size distribution up to ∼50 µm (CelluForce), and incomplete dispersion can lead to CNC aggregates of similar size, potentially enabling such bridging. However, this may introduce practical limitations, as even the 100 µm membranes were difficult to handle, and reducing the thickness to 30 µm would likely exacerbate these challenges.

Fructose is present on plant leaf surfaces and was previously shown to be bioavailable to phyllosphere bacteria [10,13,46]. In a metabolic analysis of 224 environmentally sampled leaf microbes, 76% of isolated strains were able to utilize fructose for growth [47]. Due to its biological relevance, fructose diffusion was assessed across 300 µm PDMS and PDMS hybrid membranes. The thickness of the membrane and concentration of filler polymer were chosen based on the results seen with water diffusion, where the majority of membranes showed significantly higher relative flux with a thickness of 300 µm and a filler polymer concentration of 10%. Although all hybrid membranes showed an increase in permeability to water, for a bigger hydrophilic molecule such as fructose, the measured diffusion rates were 10^2^ - 10^6^ times smaller. The enrichment of PDMS with 10% w/w Pemulen, Carbopol, or PVP, significantly increased the permeability to fructose.

Measured permeabilities were compared to isolated *Populus × canescens* leaf cuticles that were measured previously [10]. The diffusion of fructose through the 10% w/w Carbopol and 10% w/w PVP hybrid PDMS membranes was similar to that through isolated *Populus × canescens* leaf cuticles, whereas the 10% w/w Pemulen membrane exhibited higher diffusion. This indicates that fructose diffusion across biomimetic leaf-surface membranes can be tuned to match the rates observed in natural plant cuticles.

As the bulk permeability of fructose through PDMS and hybrid PDMS was biologically relevant, the survival of the phyllosphere model strain Pe299 was assessed on these membranes at 75% RH. Because Pemulen provided the greatest increase in fructose permeability in PDMS, subsequent experiments focused on PDMS containing 10% Pemulen. Most of the decline in bacterial population size occurred within the first 12 hours, after which the remaining population remained stable for at least another 12 hours (24 hours total), albeit at very low level of culturability. This represents progress relative to earlier work, in which no bacteria could be recovered after 6 hours [20]. Although both water and fructose diffusion were higher through hybrid PDMS membranes, no significant differences were observed in bacterial survival between PDMS and hybrid PDMS, or between media lacking a carbon source and those containing fructose at 75% RH, suggesting the presence of another limiting factor.

Increasing the humidity in the growth chamber from 75% to 100% RH supported substantially longer survival on PDMS membranes, with recoverable cell numbers decreasing from approximately 10^4^ to 10^3^ CFU cm^-2^ over a period of 12 days. The 75% RH condition was chosen to align with typical humidity ranges in plant growth chambers for phyllosphere research (60–90% RH) [48]. However, this setpoint usually refers to the chamber atmosphere, whereas the interior of closed plant culture boxes likely reaches saturation (100% RH) due to contributions from plant growth agar, plant respiration, and evaporation [49]. Further, processes such as transpiration and re-condensation of transpired vapor, along with the hygroscopic cuticle polymers and microtopographic water entrapment, maintain a boundary layer of air immediately above the leaf surface that is more humid than the surrounding environment [50,51]. This elevated near-surface humidity supports the formation and persistence of microscopic surface wetness in the form of thin water films and micro-scale droplets, which are critical for bacterial survival [52]. Bacteria can persist in these microhabitats even when leaves appear dry at the macroscale, because they aggregate within droplets stabilized by hygroscopic solutes and surface microstructures, particularly at moderate relative humidities where microscopic wetness can still be maintained by condensation and deliquescence [53]. Humidity is known to play a crucial role in bacterial colonization of leaves, influencing the maximum attainable bacterial load [54,55]. At lower humidities, bacterial aggregation and droplet formation can promote longer survival, which may explain the observed intermittent survival at 75% RH [52]. In contrast, the lack of chemical and structural features on PDMS may limit its ability to retain surface water and stabilize such micro-scale wet habitats and prevent active buffering of the microhabitat, further constraining survival [11,52]. Future modification to the PDMS surface with more complex chemical and structural features, may enhance water retention and further improve bacterial survival.

Notably, even at higher humidities, bacteria were not able to be recovered from membranes suspended under media lacking fructose. This result provides further validity to the observations that fructose permeability across PDMS membranes is biologically relevant and required to support bacterial survival.

The fructose-bioreporting strain *P. eucalypti* 299R FruBP_red was used to measure fructose availability to the bacteria colonizing different membranes. The bioreporter strain responds to fructose by producing unstable sYFP2[AAV] fluorescent proteins, thus reporting on currently available fructose. If fructose is degraded, the yellow fluorescent signal is degraded shortly after [13]. As expected, the bioreporter cells did not fluoresce at the time of inoculation, however, when fructose was supplied in the medium, bioreporter exhibited fluorescence four hours after inoculation, indicating the availability of fructose. Bioreporting cells were heterogeneously distributed and occurred as clusters of yellow fluorescent cells. The observed fructose diffusion at these sites results in a non-uniform nutrient distribution, with a higher fluorescent response at the center. This heterogeneity may be beneficial for replicating the diverse microenvironments present on plant leaf surfaces. Previous studies investigating availability of fructose on the surface of bean leaves and poplar cuticles detected similar outcomes: Fructose was abundant in specific sites located near veins and trichomes, while there were regions where no yellow fluorescent cells were detected [10,13]. These sites were more numerous on hybrid PDMS membranes (10% Pemulen). Survival is likely to co-occur only at the sites where diffusion of fructose and water is high. The latter would prevent bacteria from desiccation and thereby promote survival independent of the presence of fructose.

In our experiments, PDMS membranes were fixed in position for 6 days prior to inoculation, allowing growth media penetration. Membranes were then washed to remove excess surface fructose, ensuring bacterial responses reflected real-time diffusion rather than pre-accumulated sugars. Bacteria on natural leaf surfaces, however, encounter established nutrient pools that are consumed and replenished over time [46]. Therefore, future experiments using PDMS membranes as leaf proxies should omit the washing step to better mimic natural colonization conditions.

As demonstrated, reducing PDMS membrane thickness effectively increased bulk water permeability and nutrient diffusion. To further enhance nutrient transfer and bacterial survival, future work should focus on creating localized regions of higher permeability, generating nutrient-rich microenvironments akin to those on leaf surfaces. Such heterogeneity can be introduced by incorporating hydrophilic filler polymers into PDMS, either those examined here or alternatives such as polyethylene glycol or the triblock copolymer Poloxamer 188 [45,56,57]. These additives can promote hydrophilic domains within the hydrophobic polymer, improving the diffusion of water and small polar molecules. A continuous filler polymer network may form at high filler concentrations or in thinner membranes, as shown for PDMS-PVP and Carbopol-PDMS hybrids, which markedly increased the permeability of caffeine and water through 100–200 μm membranes [26,29,45]. Alternatively, targeted thinning of the PDMS surface, by introducing surface depressions on the medium-facing side or re-incorporating leaf topography, could present structural routes to generate similar nutrient-rich hotspots.

Overall, the results presented here suggest that the delivery of fructose to the surface of PDMS membranes is possible through diffusion, and that this diffusion is of biological relevance: when using membranes with a thickness of 300 µm, the diffusion of fructose is comparable to isolated leaf cuticles of walnut and poplar [10,58]. These findings were validated by the extended period of bacterial survival on PDMS membranes at 100% RH.

## Conclusion

This study advances biomimetic PDMS leaf surfaces by enriching them with filler polymers like Pemulen, Carbopol, and PVP, which significantly enhanced water and fructose permeability, matching rates observed in isolated *Populus × canescens* leaf cuticles.

Fructose diffusion proved biologically relevant, enabling extended bacterial survival of up to 12 days when incubated under high relative humidity conditions (100% RH). Bioreporter assays on 300 µm PDMS + 10% Pemulen membranes confirmed heterogeneous, leaf-like nutrient hotspots in which fructose was available. These advances overcome the previous limitations of phyllosphere related biomimetic systems, paving the way for engineered surfaces with topography and advanced fillers to support long-term microbial experiments.

## Materials and methods

### Membrane mold fabrication

A 4″ prime grade silicon wafer was used as a substrate for the membrane mold master. To prepare the wafer for the application of the dry film photoresist, it was first placed in an oven (Heratherm, Thermo Scientific) at 185 °C overnight to remove any moisture. The wafer was then removed from the oven and left to cool to room temperature. To clean the wafer, it was placed in oxygen plasma (Tergeo, PIE Scientific) at 100 W for 10 minutes.

Photolithography on negative-tone dry film resist (SUEX 300, DJ MicroLaminates) was used to produce master molds with circular wells with a diameter of 22 mm at depths of 100, 300, and 500 μm [31].

### Membrane fabrication

The membranes were fabricated as described previously [25]. Non-supplemented PDMS, from here onwards referred to as PDMS, was prepared at a ratio of 10:1 w/w base:curing agent (Sylgard 184, Dow). To prepare hybrid PDMS mixtures, the fillers Carbopol (971 P NF, Lubrizol), Pemulen (TR-2 NF, Lubrizol), cellulose microfibers (CMF, medium length; Watman), cellulose nanocrystals (CNC; CelluForce), or polyvinylpyrrolidone (PVP; Sigma-Aldrich) were first added to PDMS base and then thoroughly mixed. Following this, the PDMS curing agent was added and the mixtures were mixed again. Fillers were added at concentrations of 5%, 10%, and 15% w/w. The mixtures were degassed in a vacuum desiccator for 30 minutes to remove any air bubbles. After degassing, the mixtures were poured onto the mold, degassed briefly to ensure complete wetting of the mold, and covered with an acetate sheet (overhead projector transparency film, OfficeMax) while carefully pushing excess polymer to the side. To ensure uniform thickness, a flat glass sheet and a flat rubber sheet were used to evenly distribute an additional weight of 3.5 kg across the acetate sheet. The set-up was then cured on an 80°C hot plate (MSH-20D; Witeg) for two hours. The resulting membranes were manually peeled off the mold. To verify and record the thickness of PDMS and hybrid PDMS membranes, they were measured using a micrometer screw (Type 0806, Helios Preisser).

### Water permeability measurement

To measure the permeability of PDMS and hybrid PDMS membranes, a previously described set-up was used [10,25]. Briefly, the thickness of the PDMS and hybrid PDMS membranes was measured using a micrometer screw. Membranes or aluminium foil as a negative control (16 mm in diameter) were immobilized between a stainless-steel diffusion chamber and a stainless-steel ring. High-vacuum silicone grease was used to seal the chambers (Beckman Silicone vacuum grease, 335148). Deionized water was added to the donor chamber, and the chambers were placed in a container with silica gel overnight at 30°C, with membranes facing upwards, to remove any surface humidity. The chambers were rolled on a rolling mixer (RS-TR05, Phoenix Instrument) continuously between weight measurements. The chamber weight was recorded every day for six days using a pipette calibration balance (AND-Australia, AD-4212A-PT, 110 g/0.1 mg Pipette Calibration Balance/Accuracy Tester).

To assess the permeability of hybrid PDMS-CNC membranes, an inverted diffusion chamber setup (described below) was used to enable higher sample throughput and more efficient handling. Membranes were immobilized between the autoclaved 5 mL Eppendorf tube (filled with deionized water) and the pre-drilled cap. The assembled tubes were placed face up in a container with silica gel overnight at 30°C to remove residual moisture.

Chambers were stored inverted at 30°C between weight measurements. Chamber weight was recorded every day for six days using an analytical balance (Precisa, Series 390 Semi-Micro balance. 225 g/0.01 mg).

The measured weight loss was normalized for inherent variations in the fabricated membrane thicknesses. This normalization was performed according to equation (1):

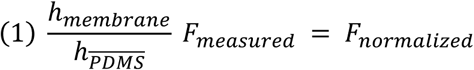

where, *h_membrane_* is the measured thickness of the membrane (μm), *h_PDMS_* is the mean thickness of the PDMS casts from the respective molds 100, 300, or 500 μm, and *F_measured_* and *F_normalized_* are the measured and the normalized weight, respectively.

Areal flux was calculated using equation (2):

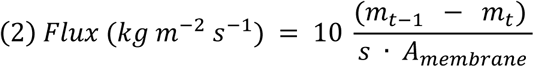

where the change in mass between time points (*m_t_*_−1_ − *m_t_*) is divided by time between sample (s) and the available diffusion area (*A_membrane_*). To further reduce variability introduced by the different experimental setups, a relative flux value was calculated by normalizing against the respective PDMS thickness control measurement. For each experiment at least five samples per material, per thickness, and per filler polymer concentration were measured.

### Construction of inverted diffusion chambers

An alternative setup was implemented following a change in laboratory environment, enabling higher sample throughput and more efficient handling. Screw caps were removed from 5 mL centrifuge tubes (Eppendorf). A 13.5 mm hole was drilled in the middle of each screw cap, and the edges of the holes were deburred using a scalpel. Afterwards, the tubes were reassembled and autoclaved. To incubate tubes membrane-side down, which simulates the abaxial side of the leaf, while a donor solution was provided from above (Supplementary Figure 1), a dedicated tube rack was designed using the software Fusion (Autodesk Fusion v2701.1.18). The racks were then prepared using the software PrusaSlicer (v2.4.2, Prusa) and printed in PLA filament (1.75 mm, Prusa) on an i3 MK3S 3D printer (Prusa) with the preset settings 0.2 mm Quality. The 5 mL centrifuge tubes were sterilized through autoclaving, while the tube rack was surface sterilized using a UV lamp (HNS 15W UV-C 254 nm, Osram) in a sterile bench for 15 minutes from every side prior to its use to minimize risk of contact contamination.

### Bacteria and culture conditions

All strains used in this study are listed in Table 1. Growth media was prepared according to the manufacturer’s recommendations. *E. coli* strains were grown on Lysogeny Broth (LB; LB broth (Luria/Miller), Roth) or Lysogeny Broth Agar (LBA; Luria Bertani Agar, Miller; HiMedia). *P. eucalypti* 299R was cultivated on Nutrient Broth (NB, Roth), Nutrient agar (NA; Roth), M9 medium or M9 agar (8.51 g L^-1^ Na_2_HPO_4_ x 2 H_2_O, 3 g L^-1^ KH_2_PO_4_, 0.5 g L^-1^ NaCl, 1 g L^-1^ NH_4_Cl, 0.49 g L^-1^ MgS0_4_, 0.01 g L^-1^ CaCl_2_, 0.2 g L^-1^ casamino acids) depending on the experiment. M9 minimal medium was supplemented with 0.4% galactose or fructose as a carbon source. Solid media (LBA, NA and M9 agar) contained 1.5% w/v agar (Agar No.1, Oxoid). Where appropriate, antibiotics were added to the media in the following concentrations: ampicillin (100 mg L^−1^), kanamycin (50 mg L^−1^) and gentamicin (20 mg L^−1^). *E. coli* strains were cultivated at 37°C and *P. eucalypti* 299R strains at 30°C.

**Table 1:** Bacterial strains used in this study and abbreviations*.

| Strain | Genetic construct | abbreviation | Antibiotic resistance | Source |
| --- | --- | --- | --- | --- |
| *E. coli* DH5α | - |  | - | Thermo Fisher Scientific |
| *E. coli* HST08 | - |  | - | Clontech |
| *E. coli* HST08 | pProbe PfruB-syfp2[AAV] |  | Amp<sup>R</sup>, Km<sup>R</sup> | This study |
| *E. coli* HST08 | pProbe_CUSPER |  | Km<sup>R</sup>, Gm<sup>R</sup> | [19] |
| *P. eucalypti* 299R | - | Pe299 | Rif<sup>R</sup> | [32] |
| *P. eucalypti* 299R | ::Tn7-MRE145 (_red) | Pe299_red | Rif<sup>R</sup>, Cm<sup>R</sup>, Amp<sup>R</sup>, Gm<sup>R</sup> | [36] |
| *P. eucalypti* 299R | pProbe_PfruB-syfp2[AAV] | Pe299FruByfp | Rif<sup>R</sup>, Amp<sup>R</sup>, Km<sup>R</sup> | This study |
| *P. eucalypti* 299R | ::Tn7-MRE145; pProbe PfruB-syfp2[AAV] | Pe299FruByfp_red | Rif<sup>R</sup>, Cm<sup>R</sup>, Amp<sup>R</sup>, Km<sup>R</sup> | This study |
| *P. eucalypti* 299R | ::Tn7-MRE145; pProbe_CUSPER | Pe299CUSPER | Rif<sup>R</sup>, Cm<sup>R</sup>, Km<sup>R</sup>, Gm<sup>R</sup> | [36] |
| *P. eucalypti* 299R | ::Tn7-MRE153 | Pe299_yfp | Rif<sup>R</sup>, Cm<sup>R</sup>, Km<sup>R</sup> | [36] |

\*Abbreviations of antibiotic resistances: ampicillin (Amp<sup>R</sup>), kanamycin (Km<sup>R</sup>), chloramphenicol (Cm<sup>R</sup>), rifampicin (Rif<sup>R</sup>) or gentamycin (Gm<sup>R</sup>).

### Plasmid construction and bacterial transformation

The plasmid pProbe_P_fruB_-syfp2[AAV] was constructed via isothermal assembly [33].

This is a reconstructed and updated plasmid of the pP_fruB_-gfp[AAV] plasmid originally constructed by Leveau and Lindow [13]. The original fluorescent protein, GFPmut3[AAV], was substituted with a newer generation fluorescent protein gene, sYFP2[AAV], encoding for a fluorescent protein with a higher fluorescent signal [34,35]. DNA fragments were obtained through polymerase-chain reaction (PCR) using Phusion High-Fidelity DNA polymerase (Thermo Scientific) following the manufacturer’s recommendations. Annealing temperatures (T_a_) were chosen based on the respective melting temperature (T_m_) of the primers (Table 2). Touchdown PCRs were performed to amplify PCR products with overlapping ends for isothermal assemblies, as described previously [36]. The P*_fruB_* promoter fragment was amplified using genomic DNA *E. coli* DH5α as a template and primers Pfrub_fw and Pfrub_rv. The yellow fluorescent protein gene (*syfp2)* was amplified from plasmid pMRE143 (Schlechter et al. 2018) using the primers ufp_fw and ufp_rv. The DNA sequence coding for the Alanin-Alanin-Valine peptides at the 3’ end of the gene were incorporated into the sequence of the primer ufp_rv to add the peptide sequence at the C terminus of protein. The β-lactamase resistance gene (ampR) and a 5’ located terminator sequence were amplified from pGRG36, a kind gift from Nancy Craig, (Addgene plasmid #16666; http://n2t.net/addgene:16666; RRID:Addgene_16666) using primers ampR_fw and ampR_rv [37]. All PCR fragments were purified using the Monarch PCR clean up kit (New England Biolabs) or the Monarch DNA Gel Extraction Kit (New England Biolabs) after separation from PCR secondary products on a 1% agarose gel following the manufacturers’ recommendations. Plasmid pFru97 [38] was digested using HindIII (Thermo Scientific) and ScaI (Thermo Scientific). The ∼5700 bp backbone fragment was gel-extracted and all fragments were then assembled into pProbe_P_fruB_-syfp2[AAV] by isothermal assembly. The assembly mix was transformed into *E. coli* HST08 competent cells (Stellar^™^, Takara Bio Inc., Japan) via heat shock, following the manufacturer’s protocol. The plasmid was then transformed into *P. eucalypti* 299R and *P. eucalypti* 299R::MRE-Tn7-145 (Schlechter et al. 2018) through electroporation (1 mm cuvette, 1.8 kV at 200 Ω and 25 µF), resulting in *P. eucalypti* 299R (pProbe_P_fruB_-syfp2[AAV]) and *P. eucalypti* 299R::Tn7-MRE145 (pProbe_P_fruB_-syfp2[AAV]) referred to as Pe299FruByfp and Pe299FruByfp_red from here onwards.

**Table 2:**
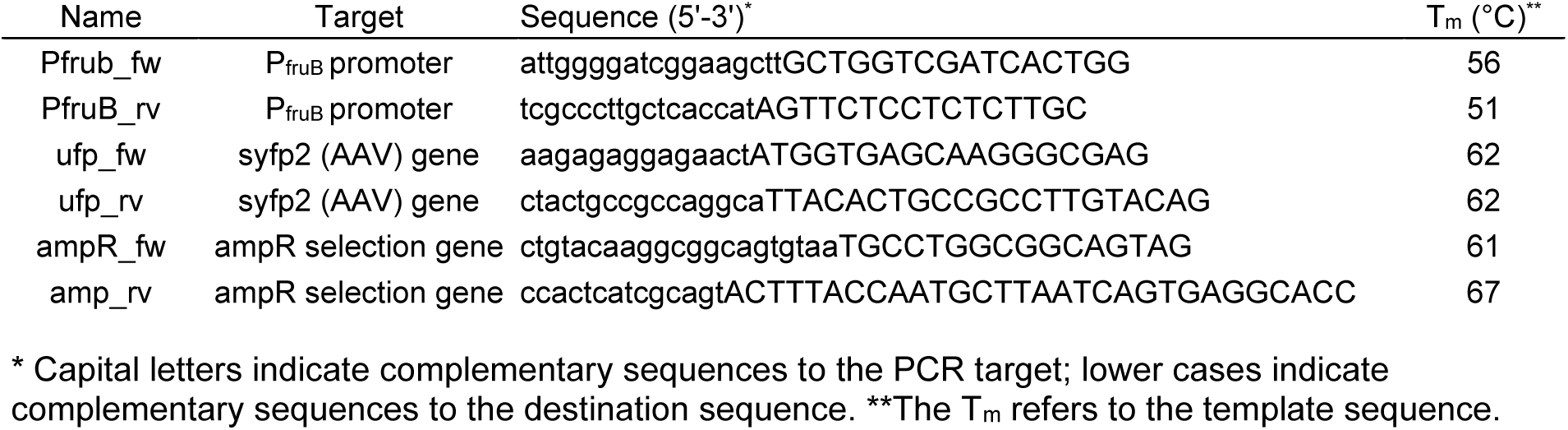
List of primers used in this study.

### Fructose diffusion measurement

To measure the permeability of the materials to fructose, diffusion chambers were used [10,39]. In short, 16 mm diameter membranes were immobilized between a stainless-steel diffusion chamber (donor chamber) and a stainless-steel ring. An additional stainless-steel chamber (acceptor chamber) was connected to the ring on the opposite side of the membrane. A 250 μm thick transparent PVC film (Office Max) was used as negative control. To further seal the chambers and to prevent leakage, the surface of the diffusion chamber components that were in contact with the membranes or each other were covered with high-vacuum silicone grease (Beckman Silicone vacuum grease, 335148) which was thinned using chloroform to allow for easier application to the chambers (Sigma Aldrich). Afterwards, both the donor and acceptor chambers were filled with 1 mL M9 media supplemented with kanamycin (50 mg L^−1^), 0.2% w/v casamino acids (Peptone, VWR Chemicals), 0.4% w/v galactose (M9_Gal, Kan_), which does not activate P_fruB_ ^[^13^]^. In addition, the donor chamber contained 3 M fructose. The chambers were then incubated while rolling on a tube mixer (RS-TR05, Phoenix Instruments) at 30°C. An aliquot of 500 μL from the acceptor chamber solution was collected after six days. This volume was replenished with fresh M9 medium supplemented with kanamycin, casamino acids, and galactose. After twelve days, the full content of the acceptor chamber (∼1 mL) was collected. All samples were stored at -20°C until the fructose concentration in the acceptor chamber was measured.

To measure diffused fructose, the fructose bioreporter Pe299FruByfp_red was used.

Yellow fluorescence was monitored with a microplate reader (CLARIOstar Plus, BMG LABTECH). For each well of a flat-bottom 96-well plate (Brand), 50 μL of an overnight Pe299FruBP_red M9_Gal, Kan_ culture, 100 μL of fresh M9_Gal, Kan_ medium, and 50 μL of acceptor-chamber sample were combined, resulting in a 1:4 dilution of the sample. When necessary, the acceptor-chamber sample was diluted further to keep the fluorescent signal from Pe299FruByfp_red within the detection range. A calibration curve with different concentrations of fructose (0, 0.025, 0.05, 0.1, 0.2, 0.39,,0.78, 1.56, 3.25 mM) was prepared within the same 96-well plate every time the assay was performed. Fluorescence was determined for 8 hours every 10 minutes. The 0 mM fructose control samples were subtracted from all samples as background correction. For the remaining concentrations, the area under the curve (AUC) was calculated using the AUC() function from the *DescTools* package in R [40]. AUC values were plotted against the fructose concentration, and a Monod model was fitted using the function nls() from the core *Stats* package, using **equation 3**:

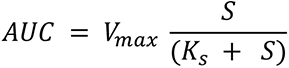

Where V_max_ is the theoretical maximum signal, and K_s_ is the specific concentration where 50% of V_max_ is reached. Rearranging this equation allows for interpolation of fructose concentrations (S) for the measured samples. Fructose diffusion values were adjusted for membrane thickness using the **equation 1** above and multiplied by the appropriate dilution factor.

### Survival of bacteria on PDMS and hybrid PDMS membranes

Prior to assembly of the inverted diffusion chambers, each side of PDMS and hybrid PDMS membranes was UV sterilized for 15 minutes. Sterile membranes were immobilized between the autoclaved 5 mL Eppendorf tube (filled with 2 mL M9 medium with kanamycin, and with or without 1 M fructose and casamino acids) and the pre-drilled cap (Supplementary Figure 2). The assembled tubes were placed in a sterile plastic box containing saturated NaCl solution to maintain a relative humidity of 75% and preincubated for 6 days at 30 °C to allow for medium to penetrate the membranes. Prior to the inoculation of bacteria, the membrane surfaces were washed twice with 600 μL sterile milliQ water to remove excess of previously permeated medium, ensuring only the impact of current fructose diffusion was captured.

Pe299_yfp was grown overnight on M9 agar plates supplemented with 0.4% fructose and kanamycin at 30°C. The bacteria were harvested and suspended in 5 mL of sterile M9 medium without casamino acids or carbon source and washed twice by centrifugation at 1150 RCF for 5 minutes at 10°C before resuspension to an OD_600_ of 0.05 (approximately 5 × 10^7^ bacteria per mL). 300 μL of bacterial suspension was sprayed over the grid of 12 exposed PDMS or PDMS + 10% Pemulen membranes twice, using an airbrush (Ultra, Harder & Steenbeck) at 1×10^5^ Pa. Pemulen was chosen as the sole filler polymer here due to the highest fructose diffusion performance (see results). The inoculation volume was chosen to be comparable to values used to inoculate *A. thaliana* plants in the Litterbox system [41].

Following spraying, four membranes were collected (1 per material and treatment) to assess Pe299_yfp population density at T_0_. Each membrane was placed in a 2 mL Eppendorf tube, filled with 1 mL PBS buffer (8 gL^-1^ NaCl (LabServ), 0.2 gL^-1^ KCl (LabServ), 1.44 gL^-1^ Na_2_HPO_4_-GPR (AnalaR), and 0.24 gL^-1^ KH_2_PO_4_ (AnalaR), pH 7.4). The samples were vortexed for 5 minutes and sonicated for 5 minutes, before colony forming units were determined on nutrient agar supplemented with kanamycin. Additional samples were incubated as above, with samples collected after 12 and 24 hours.

Survival assays were repeated under varying relative humidity (RH) conditions using pure PDMS membranes. Saturated salt solutions of NaCl, KCl, or K_2_SO_4_, plus distilled water, generated RH levels of 75%, 85%, 95%, and 100%, respectively. Long-term survival was then assessed on PDMS membranes at 100% RH, with samples collected every 24 hours over a duration of 2 weeks.

### Fructose availability assay

To assess the availability of fructose to bacteria on the surface of PDMS and hybrid PDMS membranes, the bioreporter strain Pe299FruBP_red was introduced to the surface of the membranes and then imaged. To that end, bacteria were grown to mid exponential phase in 20 mL of M9_gal, kan_ and were then diluted to an OD_600_ of 0.5 (approximately 5 × 10^8^ bacteria per mL) using fresh media. 2 × 600 μL of bacteria were sprayed onto the PDMS and PDMS + 10% Pemulen membranes mounted in the inverted diffusion setup and then incubated for 4 hours at 30°C as explained above. For imaging, membranes were dismounted from the tubes and placed directly on microscopy slides, sprayed side up.

Images were acquired without cover slip on a Zeiss AxioImager.Z2 fluorescent widefield microscope at 10 × magnification (EC Plan-Neofluar 10x/0.30 Ph1 M27 objective) equipped with Zeiss filter sets 43HE and 46HE (BP 550/25-FT 570-BP 605/70 and BP 500/25-FT 515-BP535/30, respectively), X-Cite XYLIS Broad Spectrum LED Illumination System (Excelitas Technologies), an Axiocam 712, and the software Zeiss Zen 3.5. A Z-stack was acquired for each field of view, to account for sloped sample surfaces. Red fluorescence was acquired using filter 43HE and yellow fluorescence was acquired using filter 46HE. All images were analyzed in ImageJ/FIJI Version 1.54d [42]. Maximum intensity projection of the Z-stacks were produced for each field of view.

### Data Analysis

Unless otherwise stated, all data processing, statistical analysis, and graphing were carried out in R [40]. The normality of each treatment combination was assessed using Shapiro–Wilk tests. Linear models and ANOVA were conducted using base R. Pairwise comparisons were performed using post hoc analyses with either Dunnett’s test or Tukey’s test from the *multcomp* package [43]. Plots were created using the package *ggplot2* [44].

### Data availability statement

The datasets generated and analyzed during the current study are openly available on Zenodo at: https://doi.org/10.5281/zenodo.19472079. The analysis code and scripts used in this study are available on GitHub at: https://github.com/Evan-Kear/permeable-PDMS-leaf-paper/

## Acknowledgements

The authors thank Rebecca Soffe for her contribution during the initial phase of the project and Mila Oeltjen for her assistance during data collection.

## Author contributions

E.K., M.B., V.N., and M.R.E. conceptualized the study. E.K. and M.B. performed the experiments. E.K., M.B. and M.R.E. analyzed the data. E.K., M.B. and M.R.E. wrote the first draft of the manuscript. All authors contributed to later versions of the manuscript and agreed to the final version.

## Additional Information

The authors declare no competing interests.

