## Supplementary Figures for "Developing permeable polydimethylsiloxane-based biomimetic leaf surfaces to study phyllosphere microbial ecology"

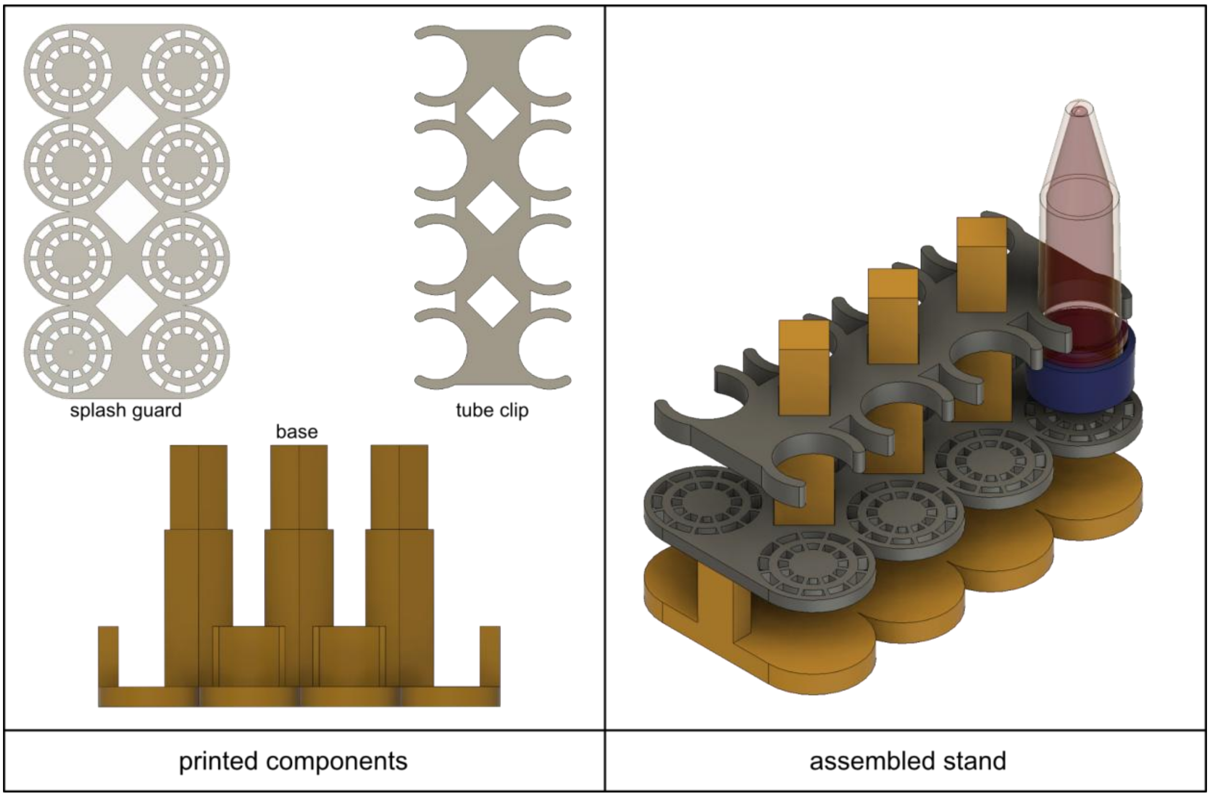


**Supplementary Figure 1. 3D printed 5 ml Eppendorf tube holder.** A tube holder was designed to allow for inverted suspension of multiple 5 ml Eppendorf tubes in Fusion (Autodesk Fusion). The components of the model (left) were sliced using PrusaSlicer (v2.4.2, Prusa) and printed on an i3 MK3S 3D printer (Prusa). The splash guard and tube clip are designed to be pressed onto the base in that order, to construct the final stand (right).

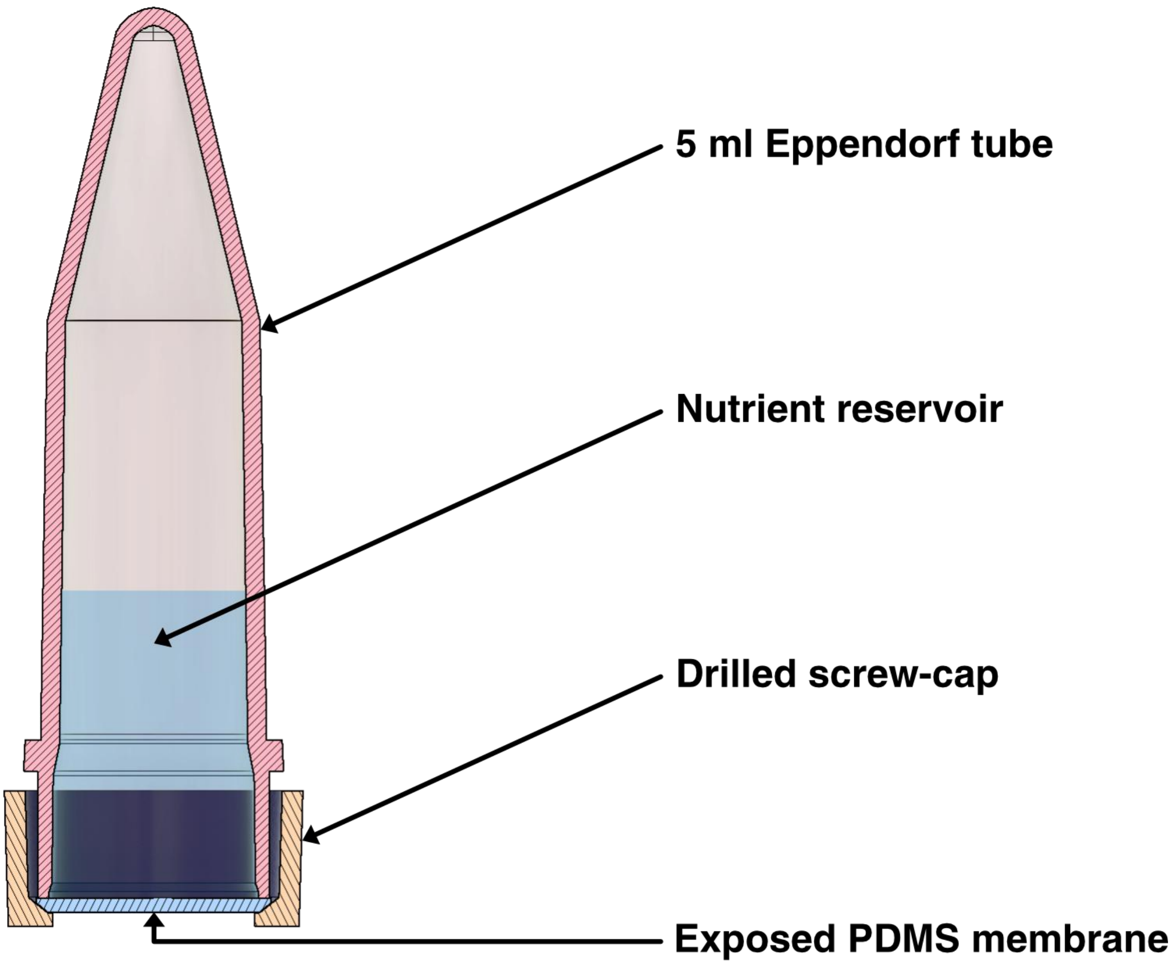


**Supplementary Figure 2. Cutaway view of the inverted 5 ml Eppendorf tube.** The inverted 5 ml Eppendorf tube acts as a reservoir for the bacterial growth medium. A PDMS, or hybrid-PDMS, membrane is fixed in place at the opening of the tube using a predrilled screw-cap. This results in the membrane being exposed to the atmosphere on the lower side, while the upper side remains in contact with the growth nutrients. This system aims to mimic the conditions present on the abaxial surface of the leaf.

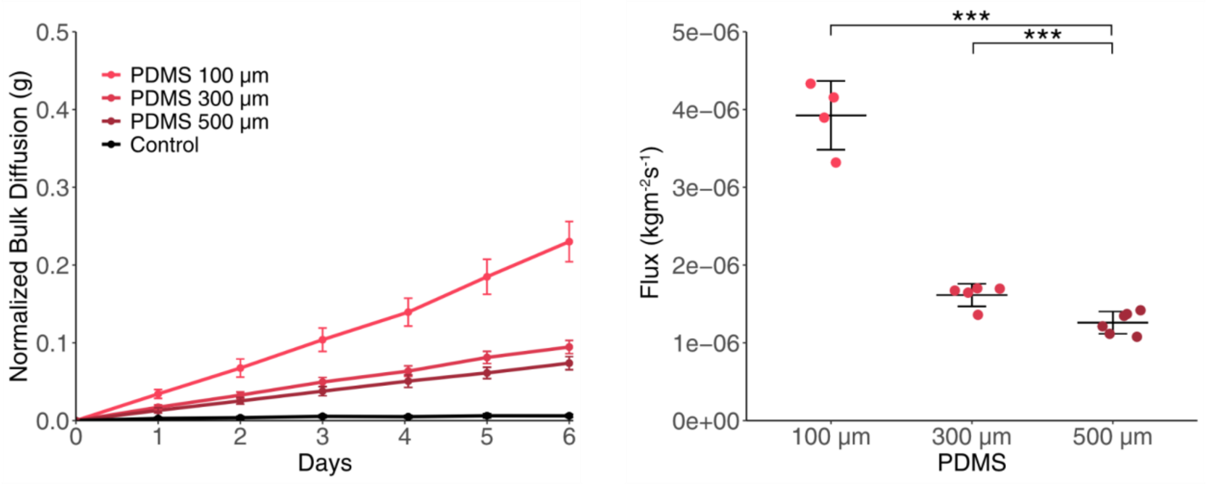


**Supplementary Figure 3: The influence of thickness on water diffusion rates through PDMS membranes.** A) Normalized bulk water diffusion coefficients through 100, 300, and 500 μm PDMS membranes. B) Absolute water flux through PDMS membranes. Asterisks denote Tukey's post-hoc p-values (p=0.05, p=0.01, p=0.001).

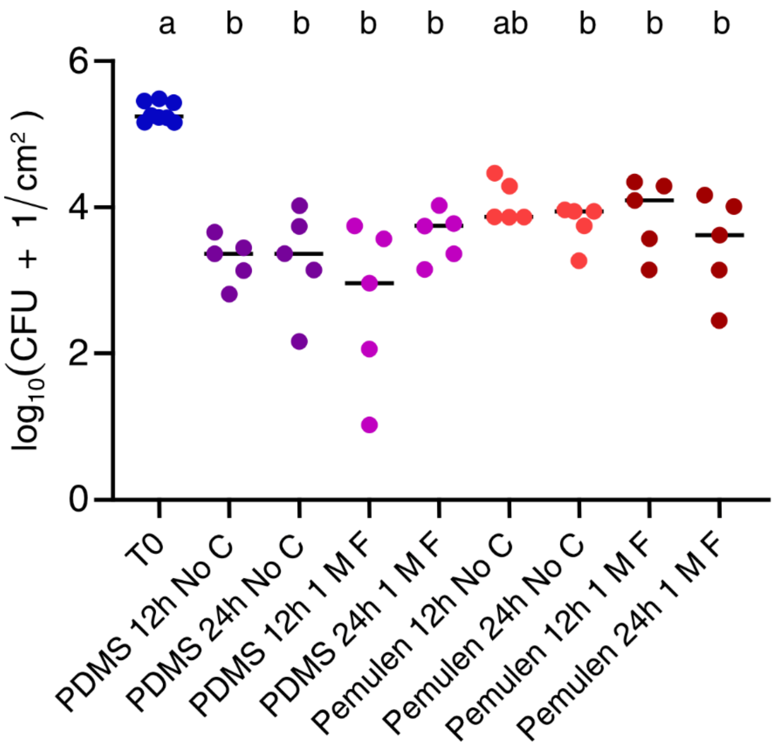


**Supplementary Figure 4. Survival of Pe299_yfp on the surface of PDMS and hybrid PDMS supplemented with 10% Pemulen.** The CFU/ml/cm$^2$ of Pe299_yfp was assessed after 12 and 24 hours. There was no significant change in survival of bacteria between treatments. No C = no carbon source in the medium and 1 M F = 1 M fructose in the medium. One-way ANOVA was performed for statistical analysis. Colony forming units at T0 were significantly different from all other samples with the exception of treatment Pemulen 12 h No C.
